# Evidence of local adaptation in Puget Sound’s threatened bull kelp (*Nereocystis luetkeana*) populations

**DOI:** 10.1101/2025.10.20.683549

**Authors:** Evelyn Abbott, Cicero Alves-Lima, Jodie Toft, Filipe Alberto

## Abstract

Stark declines in the abundance and distribution of bull kelp and reduced genetic diversity have been observed in Puget Sound, Washington, especially in southernmost areas. Consequently, introducing new variation through genetic rescue (GR) has emerged as a promising restoration strategy. However, GR success hinges on the preservation of local adaptation, which has not yet been studied in the region. To that end, we performed whole-genome sequencing on 100 gametophyte individuals from our biobank, sourced from 14 populations across Puget Sound. We identified three genetic clusters, corresponding broadly to southern Puget Sound (SPS), Whidbey Basin (WB), and the Strait of Juan de Fuca (SJF). Despite low genetic diversity and high genetic load, we found evidence of local adaptation to five environmental variables. Although salinity emerged as the strongest predictor of environmentally associated genetic variance overall, in SPS the primary drivers were average winter sea surface temperature (SST) and seasonal turbidity. We identified SNPs and enriched gene ontology (GO) terms and analyzed their allele frequencies to assess functional variation between clusters. These frequencies revealed marked differences between SJF and the other clusters, while for most genes, the highest-frequency allele in WB was also most prevalent in SPS. However, for all genes linked to winter SST and turbidity, SPS harbored distinct alleles not found in any other cluster. These findings suggest that a WB population would likely be a strong candidate donor for SPS due to its higher genetic diversity and shared adaptive variation. Further experiments are needed to validate these results and assess the suitability of WB populations for GR.

## Introduction

Genetic rescue (GR) has emerged as a method for conserving small, declining populations that require intensive conservation intervention. By introducing immigrants from a larger, more genetically diverse population, GR may increase fitness in the recipient population through a variety of mechanisms, such as heterosis, reducing genetic load, and facilitating adaptation (e.g., Hendry et al., <u>2011</u>; Tallmon et al., <u>2004</u>; Whiteley et al., <u>2015</u>). The benefits of GR have been documented in both field and laboratory experiments. GR success stories in the field have primarily centered on charismatic terrestrial megafauna, such as the Florida panther (Pimm et al., <u>2006</u>) and big horn sheep (Hogg et al., <u>2006</u>). Owing to heterosis (i.e., “hybrid vigor”), these benefits are often apparent soon after GR is implemented. Heterosis may confer a fitness advantage to F1 individuals, leading to higher survival rates in juvenile hybrid offspring (Johnson et al., <u>2010</u>) and enhancing their future reproductive success (Hogg et al., <u>2006</u>). Experiments on plants have found that compared to their pure-bred counterparts, F1 individuals may produce more viable seeds, have higher rates of germination, and increased survival rates post-germination (Finger et al., <u>2011</u>; Vergeer et al., <u>2004</u>). Studies have also found that heterosis was best predicted by the relative size and genetic diversity of the parents’ populations; it was greatest when the recipient population was small and inbred, while the donor population was large, diverse, and with low inbreeding (Pickup and Field, 2013; Vergeer et al., <u>2004</u>).

Despite its benefits, heterosis has been observed to progressively decline after the F1 generation, and outbreeding depression may arise if the donor population is maladapted to the recipient population’s local environment. Outbreeding depression has been documented in a wide variety of taxa, including plants, invertebrates, and vertebrates, with effects often emerging in the F2 generation (Edmands, <u>2007</u>). This can be due to “swamping” locally adapted alleles (e.g., Greig <u>1979</u>; Waser & Price <u>1994</u>) or epistatic interactions, such as those involving sex chromosomes, heterozygote disadvantage, or Dobzhansky-Muller incompatibilities (Fitzpatrick, <u>2008</u>). However, recent multi-generational studies provide evidence that the risk of outbreeding depression and swamping may not outweigh the potential benefits of GR. A GR field study of Trinidadian guppies found no evidence of outbreeding depression at 8–10 generations, while local adaptive variation was maintained alongside higher reproductive success, increased genetic diversity, and a tenfold rise in population size (Fitzpatrick et al., <u>2020</u>). Experimental studies on copepods have yielded similar results: although outbreeding depression was observed in the F2 generation, hybrid fitness recovered in the F3 generation and surpassed pure-bred individuals over the following 17 generations (Hwang et al., <u>2011</u>; <u>2012</u>).

Kelps of the order Laminariales are large brown macroalgae that create complex tertiary habitats in temperate and subpolar regions (Teagle et al., <u>2017</u>; Tempera et al., <u>2021</u>; Wernberg et al., <u>2019</u>) with some species in mesophotic zones of subtropical and tropical waters . Distributed across roughly 25% of the world’s coastlines, kelps are keystone species that provide habitat and food for over 1,000 species of plants and animals (Jayathilake & Costello, <u>2020</u>; NPS.gov, <u>2025</u>; Wernberg et al., <u>2019</u>). In recent decades, climate change has led to significant changes in habitat suitability, leading to the decline of 38% of studied kelp forest ecoregions (Krumhansl et al., <u>2016</u>). Many anthropogenic stressors have been linked to kelp forest decline, including ocean warming (Filbee-Dexter et al., <u>2015</u>; Smale, <u>2019</u>), eutrophication (Mason & Blain, <u>2024</u>; Moy & Christie, <u>2012</u>), sedimentation (Eckman et al., <u>1989</u>; Grossman et al., <u>2020</u>; Seapy & Littler, <u>1982</u>), and change in community composition (Bennett *et al*., <u>2015b</u>; Johnson et al., <u>2011</u>; Vergés et al., <u>2014</u>). Chronic thermal stress has led to poleward range shifts as temperatures exceed kelp physiological limitations (Sunday et al., <u>2015</u>; Weins, <u>2016</u>), while acute stress can cause widespread destruction in relatively short timeframes (Wernberg *et al*., <u>2013</u>). In some cases, this has caused hundreds of kilometers of destruction, replacing habitat-forming kelp species with turf algae (Arafeh-Dalmau *et al.,* <u>2019</u>; Smale & Wernberg, <u>2013</u>; Thomsen et al., <u>2019</u>; Wernberg et al., <u>2016</u>). Furthermore, activities such as construction, deforestation, and agriculture have led to eutrophication and sedimentation, causing reduced kelp productivity and increased competition with other algae (Moy & Christie, <u>2011</u>; Rubin et al., <u>2017</u>). These particles may also limit light availability—a process known as ocean darkening—which further decreases productivity (Blain et al., <u>2021</u>; Bass et al., <u>2023</u>; Mason & Blain, <u>2024</u>). Human activities can also destabilize marine communities, as seen in the decline of California sea otters and, more recently, sunflower sea stars due to a wasting disease (Burt et al., <u>2018</u>). The loss of these predators triggered unchecked sea urchin grazing, which in turn led to a stark decline in kelp density. In northern California, this cascade ultimately led to a major shift in the ecosystem from kelp forests to urchin barrens (Bennett & Catton, 2019; Smith & Carr, 2024).

The canopies of northeastern Pacific kelp forests are formed by two species, giant kelp (*Macrocystis pyrifera*) and bull kelp (*Nereocystis luetkeana*). These species span the range between central California and the Gulf of Alaska and have had varied regional responses to climate change (Arafeh-Dalmau et al., <u>2019</u>; Bell et al., <u>2020</u>; Cavenaugh et al., 2019; Hollarsmith et al., 2023; Starko et al., <u>2024</u>). Many of the kelp forests along the open ocean coasts of Oregon, Washington, and British Columbia have demonstrated remarkable resilience and stability, often retaining historic levels of abundance (Pfister et al., <u>2017</u>). Within this region lies the Salish Sea, an extensive fjord and estuary system that spans between Canada and the United States. It extends inland from the Strait of Juan de Fuca (SJF), then north to the Strait of Georgia and south to Puget Sound (Washington state). Like the outer coast, the western portion of SJF has largely maintained historic levels of kelp cover, forming the most robust, diverse kelp forests in the region (Pfister et al., <u>2017</u>). However, the bull kelp forests of Puget Sound are much less stable, with human activities and other stressors contributing to widespread decline (Berry et al., <u>2021</u>; Pfister et al., <u>2017</u>).

Kelp reforestation efforts are underway worldwide, with strategies aimed at both increasing kelp abundance and removing competitors. Transplantation is the most common method, which involves manually adhering sporophytes to the substrate or, more recently, deploying lab-cultured gametophytes adhered to small stones (the “green gravel” approach; ; Eger et al., <u>2022</u>; Fredriksen et al., <u>2020</u>). However, despite high success rates in other regions, many sites in Puget Sound have continued to decline. This is especially apparent in southern Puget Sound (SPS), where overall kelp cover has decreased by 63% from historic levels, with some sub-basins experiencing losses up to 96% (Berry et al., <u>2021</u>). Because bull kelp is an annual, unsuccessful transplantation sets restoration efforts back an entire year, allowing forests to decline even further. Furthermore, transplantation efforts require a tremendous amount of time, labor, and financial investment (Bayraktarov et al., <u>2016</u>; Eger et al., <u>2022</u>). Therefore, it is imperative that we develop new techniques to increase survivorship.

Recent studies have found that dwindling populations in Puget Sound are subject to strong genetic drift, which has led to low genetic diversity, high inbreeding, and the accumulation of deleterious alleles (Bemmels et al., <u>2025</u>; Gierke et al., <u>2023</u>). Coupling GR with current restoration techniques could offer a promising solution, but its potential remains limited due to insufficient knowledge on how local adaptation influences bull kelp survival in Puget Sound. However, given the steep environmental gradients across Puget Sound and the presence of many stressors, the possibility that adaptive variation exists cannot be discounted. Kelp forests near the urban centers must endure poor water quality, including toxins from urban runoff, industrial outfalls, and wastewater discharge (Washington State Dept. of Ecology, <u>2021</u>). Other human activities, such as logging and construction, have caused sedimentation in some regions, preventing holdfast attachment, smothering gametophytes, and limiting light availability (Chang & Stow, <u>1988</u>; Mumford, <u>2007</u>; Rubin et al., <u>2017</u>). Some regions—especially the Whidbey basin (WB)—experience significantly higher turbidity and lower salinity than others due to freshwater input from the Skagit and Snohomish rivers (International Joint Commission, <u>2025</u>). Additionally, limited tidal exchange in areas like small inlets and deep fjord basins exacerbates these stressors, further increasing turbidity, sedimentation, and eutrophication (Puget Sound Institute, <u>2012</u>).

To minimize the risk of swamping and outbreeding depression, it is essential to investigate local adaptation before implementing restoration strategies. In situ experiments, which gather physiological and survivorship data—such as reciprocal transplants—are ideal for gauging environmental suitability. However, transplanting and monitoring such experiments is extremely costly, labor-intensive, and difficult to permit, especially considering the sample size necessary to produce statistically significant results. Furthermore, bull kelp’s annual life cycle adds time constraints, and such an experiment could only be attempted once per year. Indirect methods, such as genomic-environment association analysis (GEA), are an effective precursor to reciprocal transplants, as they examine a broader range of populations with fewer resources.

Using these GEA results as a guide, in situ experiments could then focus their efforts on the most likely candidates for SPS GR. To that end, we sequenced 100 bull kelp gametophytes from a kelp gametophyte biobank, which were originally sourced from 14 sites ranging across SJF, WB, and SPS. This study has two main objectives: first, to identify putative genomic signatures of selection, and second, to identify candidate donors for SPS genetic rescue. Ultimately, with these genotypes preserved in our biobank, the identified candidate donors will be available for use in future GR efforts.

## Methods

### 2.1. Gametophyte sourcing, culturing, DNA extraction, and sequencing

Fertile fronds of *Nereocystis luetkeana* sporophytes were collected from 19 sites across Puget Sound in July 2019 and shipped to the University of Wisconsin-Milwaukee (Supplemental Fig. S1). Spore release was induced following the protocol described by Redmond et al. (<u>2014</u>). The resulting gametophytes were sexed based on morphology, isolated, and assigned unique codes in the UW-Milwaukee germplasm bank.

Vegetative propagation was conducted in 25 cm² polystyrene vented culture flasks (Nunc) containing 20 ml of sterile Provasoli-enriched semi-artificial seawater (Instant Ocean Salt) with a salinity of 34 and pH 8.2. Cultures were maintained under 40 μmol photons·m⁻²·s⁻¹ of irradiance from full-spectrum white fluorescent bulbs covered with red cellophane. To promote vegetative growth, the biomass was fragmented weekly using a sterile pestle.

DNA was extracted from 102 male and female gametophyte tissue samples (see Supplemental Table S1 for sample details and coordinates). A total of 30–90 mg of tissue-cultured gametophyte tufts were sampled from T75 (20 ml) flasks using wide-bore 1 ml tips. Excess moisture was removed by centrifugation at 14,000 RPM for 2 min at 4°C using silica spin columns (Macherey-Nagel) before weighing on an analytical scale.

For tissue homogenization, the tissue was transferred to 2 ml U-bottom tubes with two 5 mm tungsten carbide beads, flash-frozen in liquid nitrogen, and ground three times in a QIAGEN TissueLyser at 25 Hz for 1 min per cycle. The racks were cooled on dry ice for 5 min between rounds. The resulting fine powder was resuspended in the extraction buffer from the Macherey-Nagel Plant and Fungi DNA Kit (PN: 740120.50) and incubated at 25°C for 40 min with vortexing at 1,400 RPM. All subsequent steps followed the manufacturer’s protocol.

To reduce polysaccharide contamination, a final purification step was performed using the Omega MagBind DNA/RNA Kit (PN: M6932-00S). DNA quality and integrity were assessed using NanoDrop spectrophotometry, Qubit fluorimetry (DNA Broad Range Kit), and 1.5% agarose gel electrophoresis. All samples had A260/A280 and A260/A230 ratios above 1.8, with yields exceeding 1 µg. Finally, library preparation and whole-genome sequencing were conducted by the University of Wisconsin Milwaukee Biotechnology Center.

### 2.2. Sequence data processing and genotyping

Detailed descriptions of the data processing pipeline can be found on Github (https://github.com/evelynabbott/bullkelp_WGS). The annotated *N. luetkeana* reference genome (Alves-Lima et al., <u>2025</u>) was indexed using bowtie with --local option. Adapter trimming was conducted using cutadapt (Martin, <u>2011</u>), and the resulting files were sorted and indexed into BAM files using samtools (Li et al., <u>2009</u>). VCF files were generated using the bcftools toolkit mpileup option and called with the --ploidy 1 option, yielding a total of 2,468,726 SNPs.

Sequencing depth and site quality were assessed using vcftools. Coverage ranged from 1.001 to 16.403, with a mean coverage of 10.367 (Supplemental Fig. S2a). The fraction of missing sites per sample ranged from 0.036 to 0.444, with a mean of 0.0623. Site quality ranged from 3.010 to 999, with a mean quality of 584.904. The final VCF file was generated using vcftools, with filters set to a minimum depth of 10, minor allele frequency greater than 0.045, max proportion of missing data 0.9, and minimum quality 30. This removed 35 samples from the dataset, yielding a total of 65 samples for the remainder of this study. This VCF file was then saved as a genotype matrix and exported into R.

### 2.3. Population structure

Genetic distances and multidimensional scaling (MDS) were calculated using the R package vegan functions rda() and cordist(), respectively, using the genotype matrix as input. Identity-by-state (IBS) analysis was performed using the ANGSD (Korneliussen et al., <u>2014</u>) --doIBS function, with a genotyping rate cutoff of 0.5 and a minimum individual count of 64. Admixture analysis was conducted using NGSadmix version 3.2 (Skotte et al., <u>2013</u>), using the beagle file output from ANGSD as input. Linkage disequilibrium (LD) was calculated for each population separately, so the VCF file was split into separate files, each containing individuals belonging to a specific admix cluster. Pairwise LD was calculated across the 34 largest scaffolds using plink v1.9.0-b.8 (Chang et al, <u>2015</u>), with a minimum distance of 100 SNPs (--ld-window option). Due to the high number of SNPs, an additional filter was used to target LD between SNPs within 1000 kb windows (--ld-window-kb option). LD decay was assessed by calculating the average r^2^ values associated with binned distances between sites (Fig. 1c). Because we used haploid specimens, heterozygosity was estimated at the site level as the proportion of SNP loci in the filtered VCF that were heterozygous among the individuals sampled from each site.

**Figure 1.**
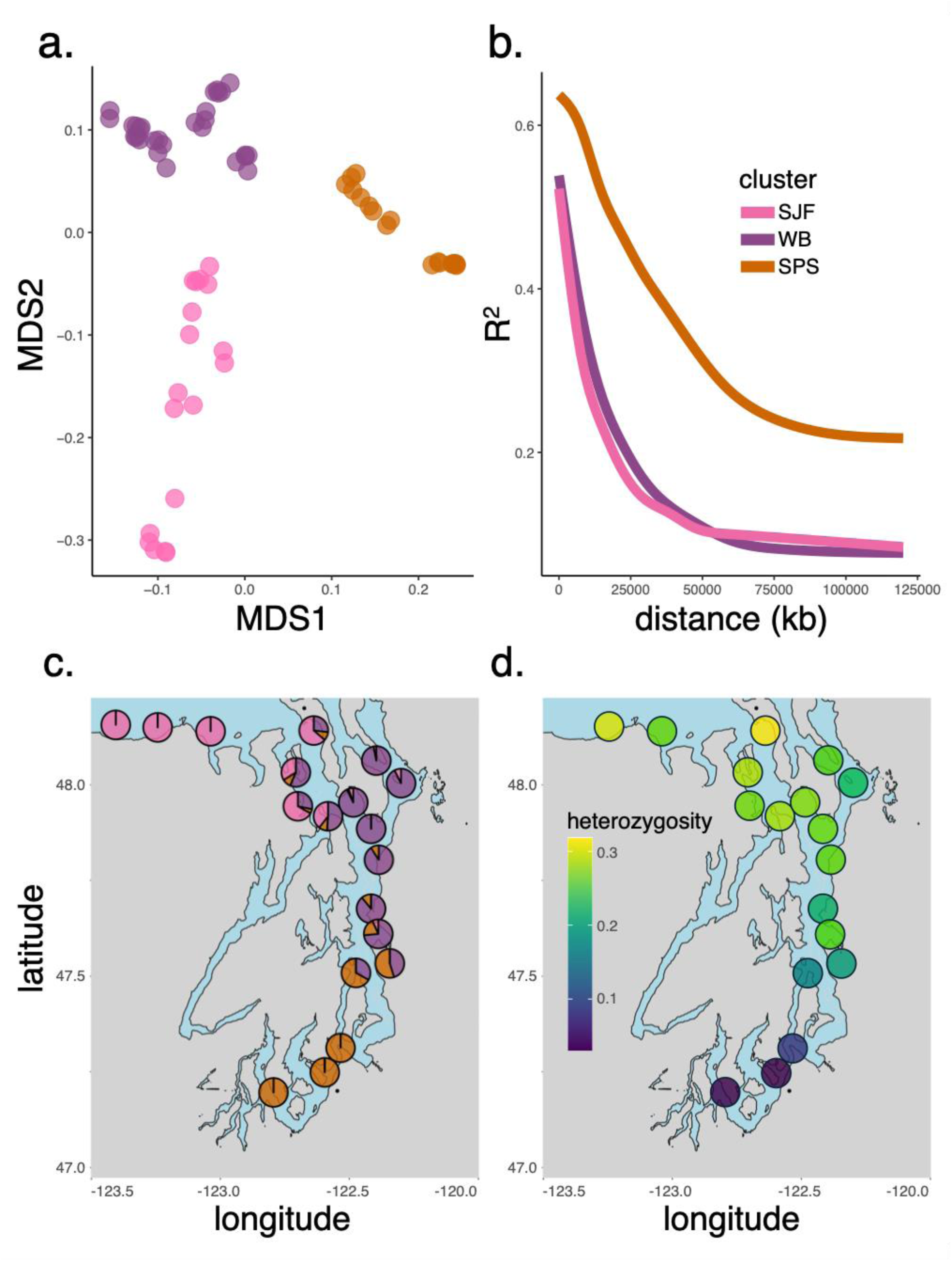
Three genetic clusters across three populations of *Nereocystis luetkeana* at Puget Sound: South Puget Sound (SPS; organge), Whidbey Basin (WB; purple), and the Strait of Juan de Fuca (SJF; pink) **a.** Multidimensional scaling (MDS) showing pairwise distances between samples. Each point represents a sample, colored according to admixture cluster. **b.** Linkage disequilibrium decay in the three admixture clusters. The points represent pairs of loci, and the axes correspond to their distances in kilobases (x-axis) and significance (y-axis). **c**. Admixture of each site, with colors corresponding to genetic clusters. **d**. Heterozygosity of each site, with the highest heterozygosity in yellow (0.32) and the lowest in purple (0.03).

### 2.4. Sourcing environmental data

Environmental data were sourced from the National Oceanic and Atmospheric Administration (NOAA) CoastWatch database (coastwatch.noaa.gov). In total, seven environmental variables were included in this study, corresponding broadly to sea surface temperature (SST), salinity, light availability (PAR), and turbidity (kd490). Two types of environmental data were used: 1) fine-scale, site-specific measurements, and 2) broad scale measurements covering the entire region. Site-specific measurements for each variable were sourced from buoy data, and the exact values are detailed in the supplemental materials (Supplemental Table S1). To avoid potential biases due to anomalous events (such as extreme storms or heatwaves, which do not typically occur), data ranging from 2012 to 2023 were included and averaged seasonally.

In cases of missing data in raster files, values were interpolated using the R package raster. A circular focal weight matrix (“moving window”) with a radius of one cell was generated for each raster using the function focalWeight(). The focal() function with the “mean” and “na.rm” options was then used to fill the missing data, using the rasters and weight matrices as input. Lastly, cell size was standardized across all rasters using the function disaggregate() with the “bilinear” method option, which locally interpolates the values of the new cells.

Notably, only variables that were consistently measured at every one of our sites could be included in this study (seasonal SST, PAR, and kd490). Although the CoastWatch database salinity measurements did not encompass the SPS, including this variable was deemed essential due to the substantial spatial variability indicated by buoy measurements. Thus, salinity data was sourced from the Salish Sea Model (SSM, Yang and Khangaonkar, <u>2010</u>), a predictive coastal ocean model for restoration planning, estuarine research, water-quality management, and climate change response assessment. The model is calibrated using quarterly observations by the Department of Fisheries and Oceans (Canada), and it takes into account currents and freshwater inflows. Using a custom Python script, salinity data was extracted from 7,219 SSM nodes, averaging across four daily time points per month in 2014. The resulting data was then imported into ArcGIS and interpolated using the spline with barriers function to prevent interpolation across land. The resulting raster was then exported to R for downstream analyses.

### 2.5. RDA-forest analysis

Due to geographic isolation in SPS, we suspected that regression models would not be sufficient for assessing drivers of natural selection in our dataset. To circumvent this, we used RDA-forest, which employs Random (RF) and Gradient Forest (GF) methods to separate genetic variation due to geographic location from variation due to environmental factors. RF is a machine learning technique that has a distinct advantage over linear regression in cases where the response variables and predictors are non-linear and/or involve multi-way interactions between predictors (Calle et al., <u>2011</u>; Ogutu et al., <u>2011</u>; Sarkar et al., <u>2015</u>). It uses decision trees to compute the importance of each predictor (in our case, environmental variables). However, applying the RF algorithm to genomic data presents many challenges due to the large number of response variables (individual SNPs). GF (Ellis et al., <u>2012</u>) is an adaptation of RF, designed to analyze a very large number of response variables. It does this by running RF on each response variable and reporting importance as the average portion of variation (R^2^) explained over all response variables.

Although GF can analyze far more response variables than random forest, it is still not ideal for datasets with thousands of SNPs. Thus, we used the R package, RDA-forest (github.com/z0on/RDA-forest), to summarize the data, representing the most significant principal components (PCs) calculated from genetic distances using our genotype matrix as input (Matz & Black, 2024). RDA-forest relies on the *mtry* criterion to determine which predictor in a group of correlated predictors is the true driver. At a high *mtry*, true predictors will rise in importance while the unimportant ones will not. An additional selection is performed by evaluating relative predictor importance: predictors whose importance is less than a certain fraction of the most significant predictor (set to 0.1) are discarded.

We ran RDA-forest, including the first five PCs with our environmental variables as predictors, and ran 50 iterations. To account for the effect of geographic isolation and distance, we included the pairwise marine distances between our sites, which were adjusted using the RDAforest gcd.dist() function. To eliminate unimportant predictors that are nevertheless correlated with important ones, we used the two *mtry* selection criteria listed above, with a threshold of 0.3 for the proportion of positive replicates. SST August and PAR winter did not pass the *mtry* selection criteria and were removed from the rest of the analysis, leaving salinity, PAR summer, kd490 spring, kd490 winter, and SST winter. PCA and adaptive neighborhood maps were plotted using the RDA-forest function plot_adaptation() with rasters of environmental variables (from NOAA CoastWatch and SSM) and predicted environmentally driven genetic variance as input (Fig. 3a,b).

### 2.6. Latent factor mixed models

Using the R package lfmm, latent factor mixed models (LFMM) were used to identify SNPs associated with the environmental variables selected by the RDA-forest *mtry* criterion (salinity, PAR summer, kd490 spring, kd490 winter, and SST winter). LFMMs are statistical regression models that test associations between a set of response variables (genotypes) and explanatory ones (environmental data). These models incorporate latent factors (unobserved variables) that account for confounding effects resulting from population structure. Because the R package lfmm does not allow missing data, imputation was performed on the genotype matrix (generated from the VCF file described in Section 2.3). To account for variation and isolation between admix clusters, the three clusters were imputed separately, with missing alleles filled using the most common allele within each cluster. The inputs for this analysis included the imputed genotype matrix and environmental parameters for each site. Modeling was conducted using the function lfmm_ridge(), and p-values were calculated using lfmm_test(). Finally, to minimize false positives, q-values for each SNP were calculated using the qvalue() function from the R package qvalue.

### 2.7. TopGO functional analyses

Functional analysis of SNPs correlated with environmental variables was conducted using the R package topGO. SNPs identified through LFMM were used as input, with a significance cut-off of q ≤ 0.01. Biological process (BP), cellular component (CC), and molecular function (MF) gene ontology (GO) terms matching the input SNP data were identified using the methods package new() function. Gene IDs and their corresponding GO terms were sourced from our annotated reference genome, which is publicly available through the Joint Genome Institute (JGI) Phycocosm portal (Grigoriev et al., <u>2021</u>). The annotation includes 21,728 annotated genes and 7,113 total GO terms. Enrichment analysis was conducted using the topGO runTest() function, with the elim algorithm and Fisher statistic settings . Finally, summary tables were generated using the topGO function GenTable(), with “topNodes” set to 365 (BP, MF) and 303 (CC).

We also examined the SNPs identified by LFMM, which were associated with significant terms identified by topGO. In other words, we considered the significant SNPs (q ≤ 0.01) located within genes whose annotations matched those of the significant GO terms. Using the input matrix for LFMM (generated from the VCF file), we assessed the genotypes of each individual at each SNP. In cases where only a single SNP was identified within a gene, it was treated independently, whereas multiple SNPs within the same gene were grouped and analyzed as haplotypes . Most gene annotations were associated with only a single SNP identified by LFMM, though two were associated with more (GTPase activity, D-arabinono-1,4-lactone oxidase activity). In the latter case, all sequences are displayed . Additionally, two highly significant terms were identified that did not match annotations of identified genes (Supplemental Table S2; P-type proton-exporting transporter activity and anion binding). This is due to the parentChild algorithm used by topGO (Grossman et al., 2007), which returns a parent term if it is shared by multiple genes. In these cases, the next most-significant genes with direct annotations were visualized. Documentation including all GO terms and associated SNPs can be found in the supplemental materials (Supplemental Table S2).

### 2.8. SnpEff

SnpEff is an open-source, variant annotation and prediction tool that uses an annotated reference along with SNP data (VCF file) to predict the effect of genetic variants on gene function (Cingolani, <u>2012</u>). Because bull kelp is not a model organism, a custom SnpEff database was built using the reference genome (Alves-Lima et al., <u>2025</u>) and tab-delimited annotations as input. The VCF file was used as input to the database, and SnpEff used interval forest predictions to search for and then annotate these variants as intronic, untranslated regions, upstream, downstream, splice sites, or intergenic regions. Following this, it identified synonymous and non-synonymous mutations concerning amino acid replacement, start or stop codon gains or losses, and frameshifts. Finally, SnpEff returned an output table for each sample listing the predictions for each variant, including chromosome, position, genic region, and its expected impact on gene function: sorted by “high,” “moderate,” and “low” impact. For further analysis, these files were imported into R and concatenated, allowing for analysis by source population and variant impact level.

## 3. Results

### 3.1. Population structure

NGSadmix revealed three primary genetic clusters, each predominantly found in a specific region: SJF, WB, and SPS (Fig. 1a). Individuals from the three western-most regions of SJF belonged only to a single cluster, which was not detected south of WB (Fig. 1c. The individuals from the three southernmost sites in SPS also belonged only to a single cluster; however, unlike the SJF cluster, this one was detected at nine different sites throughout the region, including at low frequencies around the western side of WB.

LD decay was substantially diminished in the SPS compared to the other two clusters, which exhibited little difference (Fig. 1b). Heterozygosity levels across the region largely supported the results of LD decay. Population heterozygosity largely corresponded to proximity to the ocean, with the highest values detected at Admiralty Head in SJF and the lowest at Day Island in SPS (Fig 1d; 0.32 and 0.03, respectively). SPS sites had significantly reduced heterozygosity compared to those in WB (p = 0.0004) and SFJ (p = 0.0002), with a mean heterozygosity of 0.10. Although the mean heterozygosity of SJF and WB was not as disparate as compared to SPS (0.29 in SJF and 0.26 in WB), this difference was still significant (p = 0.01).

### 3.2. Environmentally-associated genetic variation

#### 3.2.1 The relative importance of environmental variables on genetic variance

RDAforest was used to identify the major environmental predictors of genetic divergence across the region. Using predictors retained after mtry-based selection, the proportion of total genetic variation explained by the model was 0.23. Of the five retained predictors, salinity was the most important, explaining approximately one-fifth of the predicted variation (Fig. 2a). Turnover curves reveal the thresholds at which genetic transitions occur in association with environmental changes: large genetic transitions appear as distinct steps in the curve. Such transitions were most apparent in the SST winter and kd490 winter curves (Fig. 2c,e). For SST winter, a large step was observed at around 8°C, capturing more than half of the associated genetic variance (Fig 2c). Kd490 winter had a similarly strong transition, with a step at 1.5 kd490 which captured roughly one-third of associated variance (Fig 2e). Comparatively, the salinity, PAR summer, and kd490 spring transitions were more gradual, although salinity and kd490 winter had steps at 28ppt and 46.7, respectively (Fig 2b,d,f).

**Figure 2.**
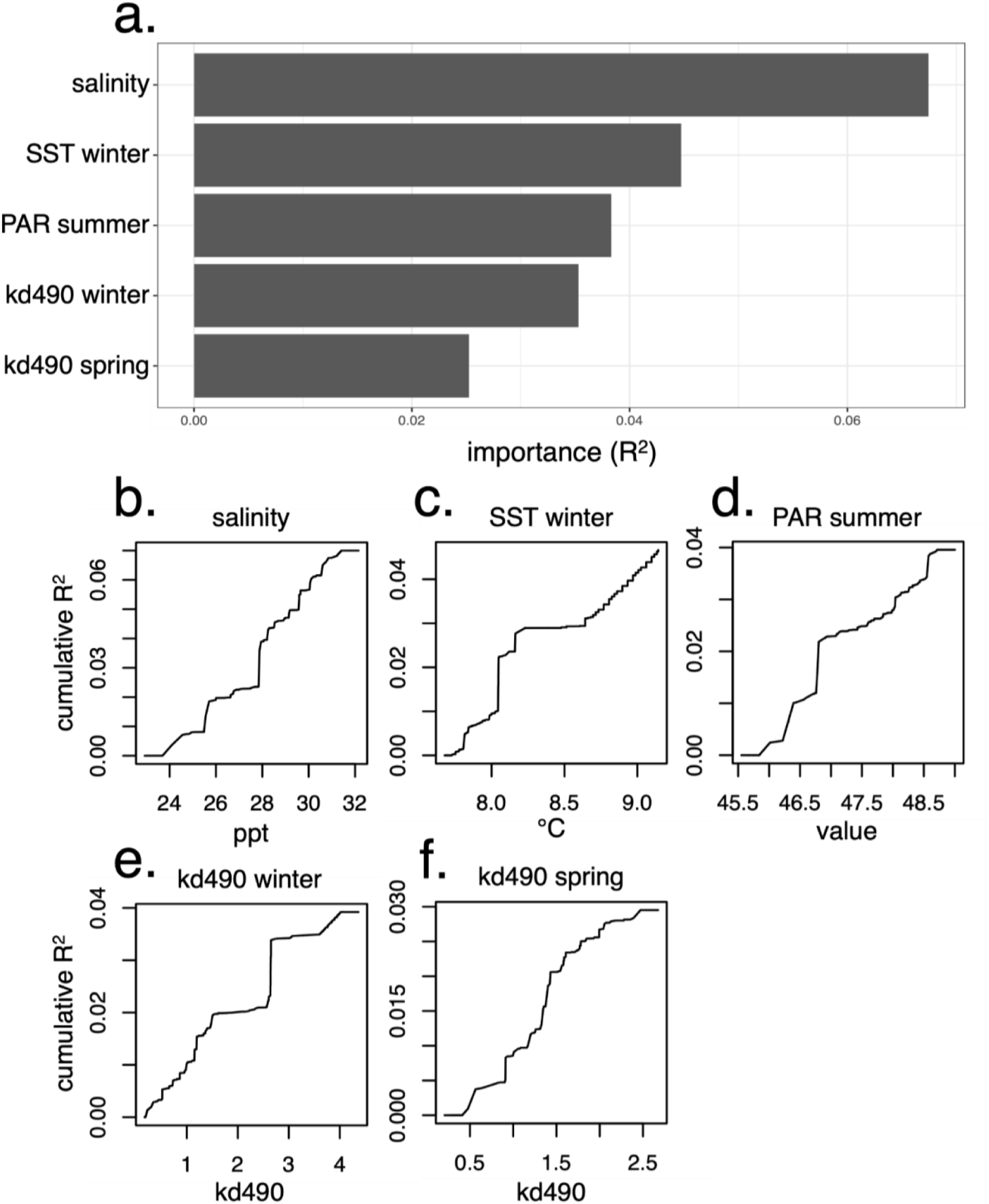
Proportion of genetic variation explained by *mtry*-retained predictors in the RDAforest model in *Nereocystis luetkeana* from Puget Sound. **a**. Bar chart showing the proportion of genetic variance explained by each predictor (x-axis and y-axis, respectively). **b-f**. Turnover curves showing cumulative genetic variation captured across each predictor range (**b**, salinity; **c**, PAR summer; **d**, kd490 spring; **e**, kd490 winter; **f**, SST winter).

A geographic map colored according to the first three principal axes of predicted genetic PCs was generated to visualize putative adaptation across the landscape. The predicted “adaptive neighborhoods” showed marked differentiation between the western-most region of SJF (green-browns) and the rest of Puget Sound (Fig. 3a, indicated by highly contrasting colors). Between the green-brown neighborhood and the western side of Whidbey Island (hot pink), RDAforest detected a small adaptive neighborhood (silver-green) that encompassed three of our sampling sites (PT, KR, FB; Supplemental Fig. S1), a boundary that was positively correlated with salinity and negatively correlated with kd490 spring and winter (Fig. 3b). Despite the short 5.3km distance between the nearest sites on either side of the boundary (FB and DB; Supplemental Fig. S1), this neighborhood was ostensibly distinct from the southern and western sides of Whidbey Island. Though still differentiated, the rest of the neighborhoods were comparatively less distinct from one another. The region occupied by the WB genetic cluster formed a single neighborhood (pink), which was negatively correlated with both PAR summer and SST winter. The purple neighborhood encompassed most sites in SPS (Fig. 3a; purple, LP, PV, SB, DI; Supplemental Fig. S1, which was correlated with both kd490 variables and, to a lesser extent, SST winter (Fig. 3b). The blue neighborhood comprised a single site (Fig. 3a; SQ) and was correlated with SST winter (Fig. 3b).

**Figure 3.**
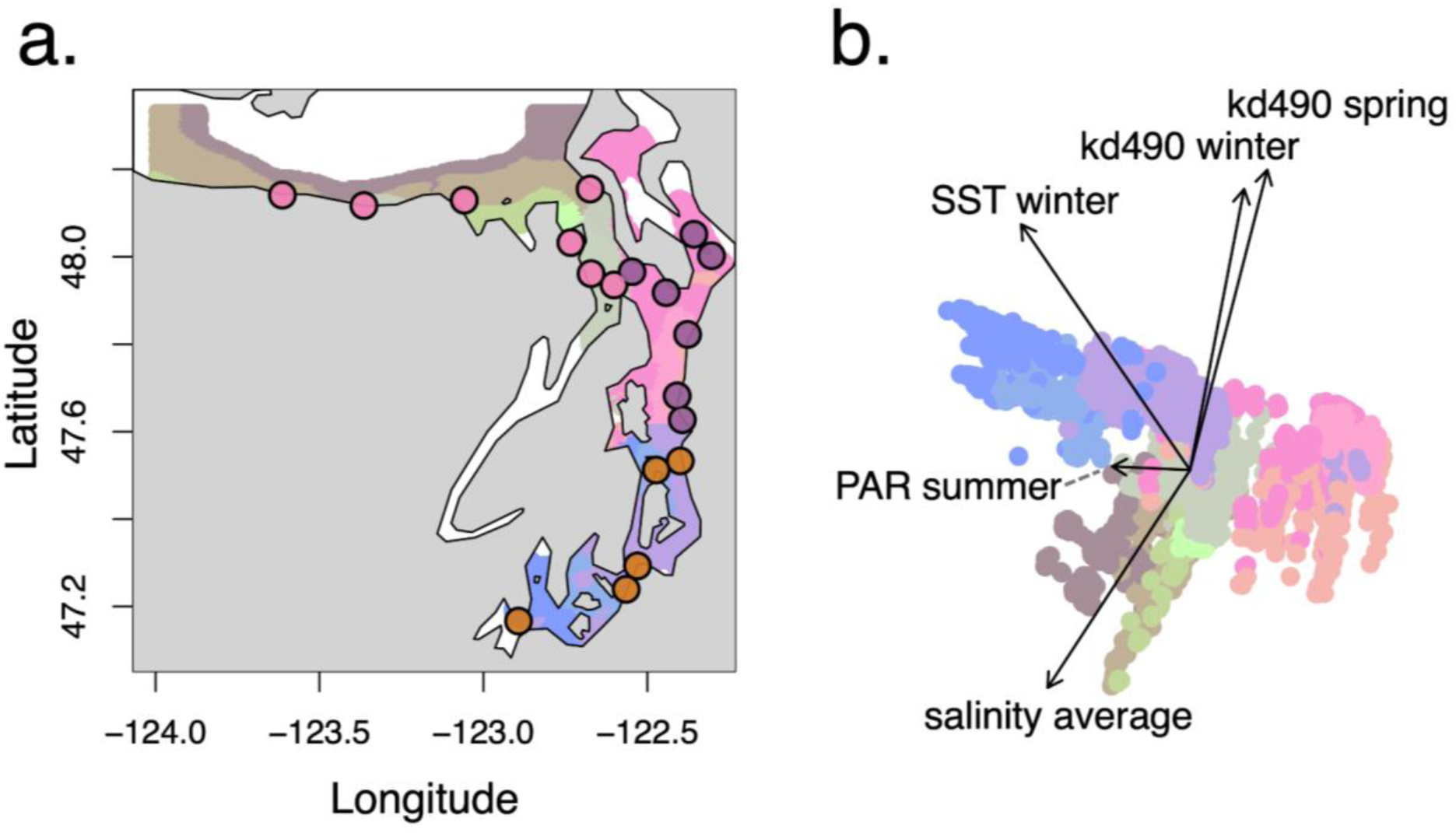
Environmentally associated genetic variation in *Nereocystis luetkeana* across Puget Sound . The colors (**a.** landscape colors, **b.** point colors) correspond to predicted genetic principal component (PC) scores along the first three principal axes of environment-associated genetic variation. Similar colors suggest similar genetics, while contrasting colors suggest differential genetics associated with the corresponding environmental variation. **a.** Map showing sites sampled in this study (points) colored according to genetic cluster (orange, SPS; purple, WB; pink, SJF). Points falling on the same map color suggest individuals at those sites may share adaptive genetic variation. **b.** PCA showing the first two principal component axes of RF predictors across the landscape, with points corresponding to locations across the region. Arrows denote the directions of the five important environmental gradients, aligned with the predicted principal axes via linear regression.

#### 3.2.2. SNPs associated with environmental variables

LFMM identified 14,576 SNPs (p ≤ 0.05) significantly associated with the environmental variables identified as important predictors by RDAforest. Despite ranking as the most important, salinity was not the predictor associated with the greatest number of SNPs (141). Instead, SST winter was associated with the greatest number by far (12,249) followed by PAR summer (1,725), and kd490 winter (413). Kd490 spring was associated with the fewest number of SNPs, accounting for merely 0.3% of all environmentally-associated SNPs (48 SNPs). Apart from the kd490 spring SNPs, which were concentrated at scaffold 25, environmentally associated SNPs were widely distributed throughout the genome, encompassing 97 of the 1,561 total scaffolds. Overall, the 31 scaffolds that contained the greatest number of SNPs accounted for the majority of those captured by LFMM (13,203 SNPs, 90.58% of the total), with scaffold 2 containing the largest number (1,002 SNPs, 8.72% of the total). Despite being less than half the size of both scaffold one and two, scaffold 28 contained the second greatest number of SNPs (911 SNPs, 7.93% of total). Of the 97 scaffolds, 55 hosted fewer than 10 SNPs, and 35 of these hosted only a single SNP.

In total, 37 GO terms were enriched from genes identified by LFMM (Fig. 4; p ≤ 0.01). PAR summer was associated with the greatest number of terms (13), followed closely by SST winter (eleven), while kd490 winter was associated with the fewest (2; Fig. 4b,c,e, respectively). Some environmental variables were associated with groups of highly similar GO terms. SST winter SNPs were enriched with five terms linked to pyrimidine biosynthesis and four linked to helicase functions (Fig. 4b). PAR summer was enriched with six terms related to motor protein functions, three related to trehalose, and three related to methionine (Fig. 4b). The two kd490 variables shared one term (inositol oxygenase activity), and kd490 spring had two terms related to nitric oxide (Fig. 4d,e).

**Figure 4.**
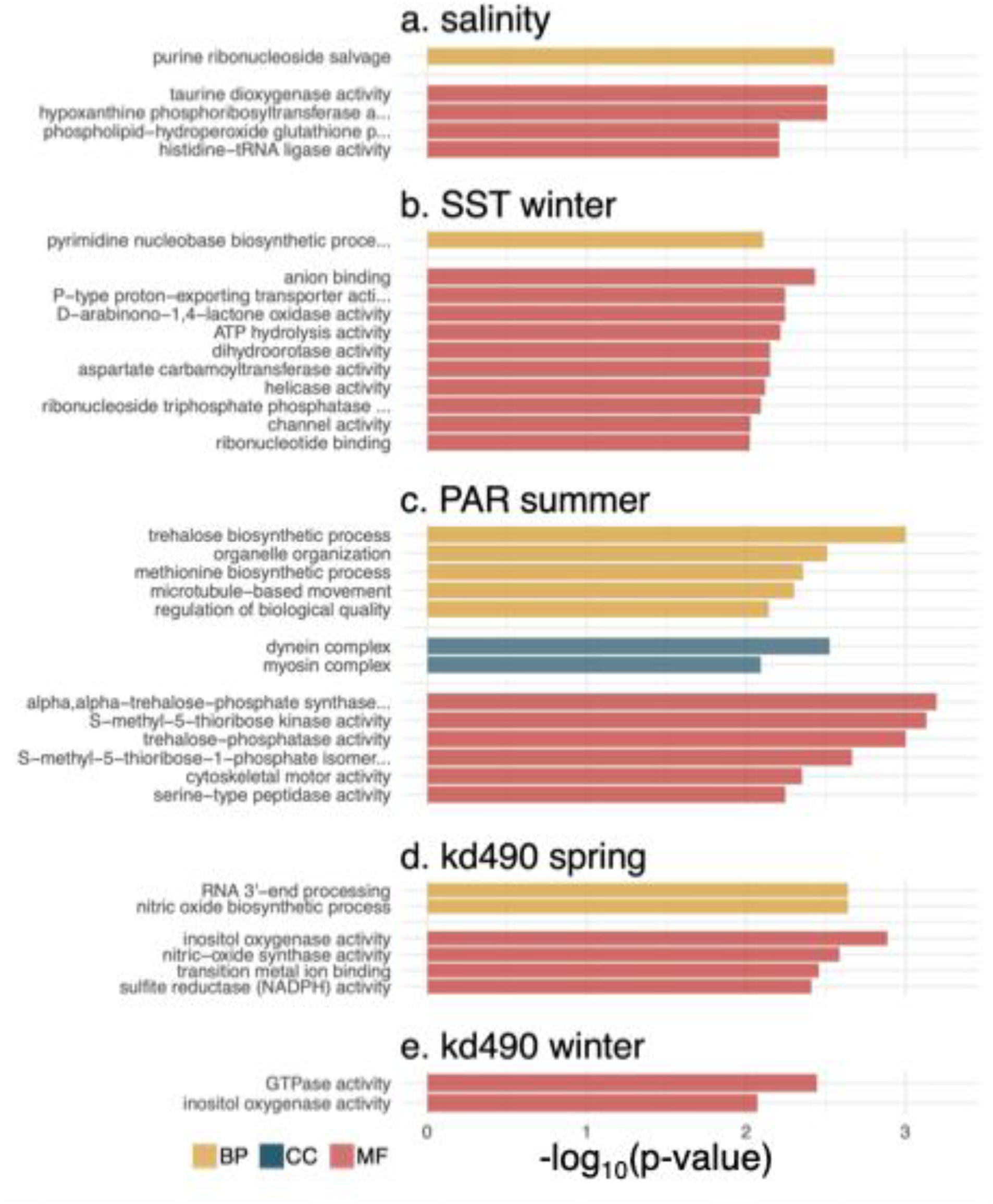
GO terms associated with environmental variables of interest (top to bottom: salinity, PAR summer, kd490 spring, kd490 winter, and SST winter) in *Nereocystis luetkeana* at Puget Sound. The level of enrichment (-log_10_(p-value)) and GO terms are displayed on the x and y axes, respectively. Colors correspond to biological process (orange), cellular component (blue), and molecular function (red). Only significantly enriched (pvalue < 0.01) GO terms are plotted.

To assess functional variation across the landscape, we analyzed the allele frequencies of SNPs identified by LFMM whose annotations matched the most significant GO terms returned in the previous analysis. The b2 PAR gene was associated with six haplotypes, the greatest number of unique haplotypes (Fig. 5b2;). Four of these haplotypes were present in SJF and all were present in WB, though only one was present among SPS individuals (Fig. 5b2; red). Differences between SJF and the other sites were most apparent among the salinity-associated haplotypes (Fig. 5a).

**Figure 5.**
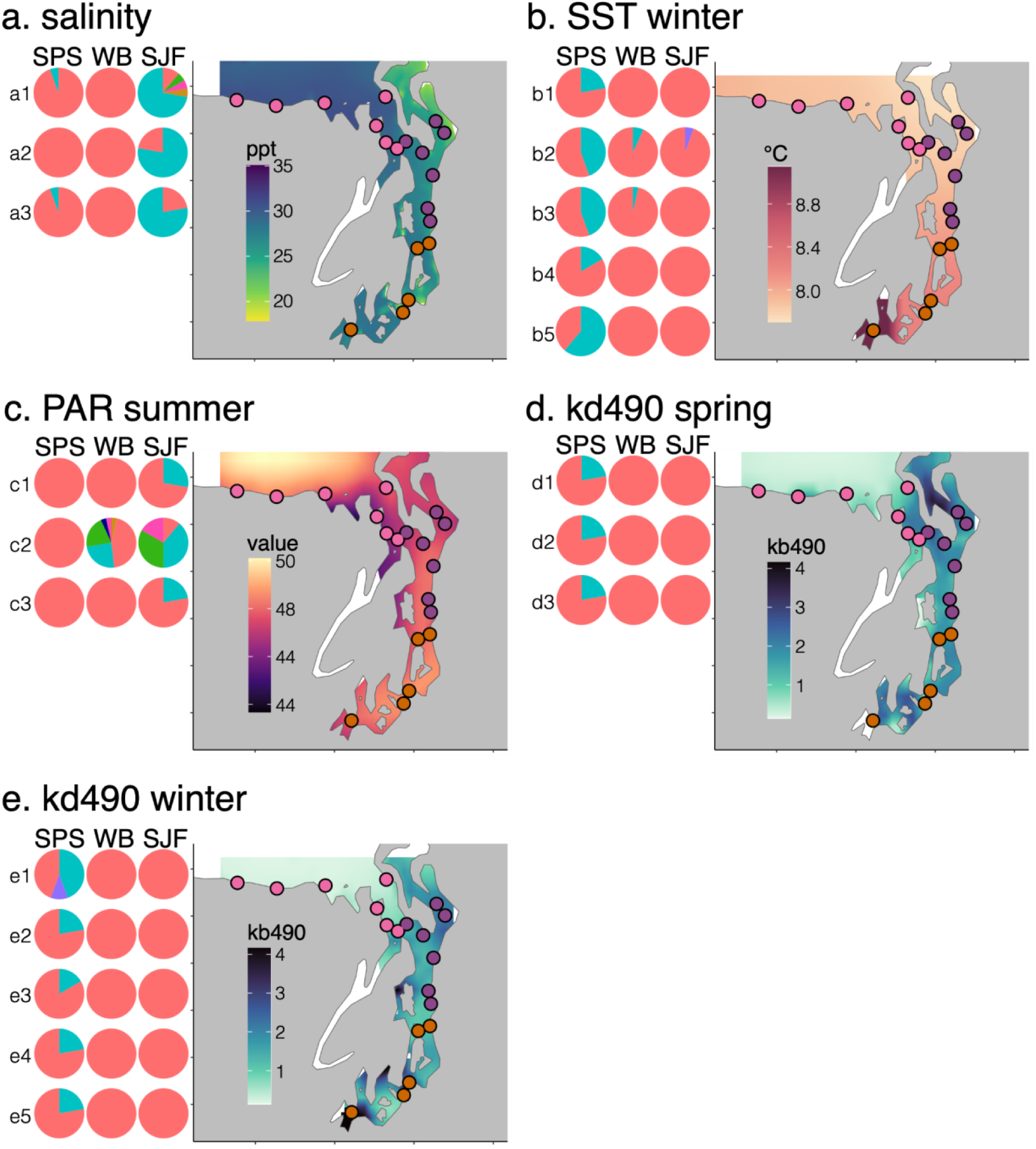
Frequencies of haplotypes associated with significant GO terms (left) and maps of environmental variables across the landscape (right) for salinity (**a**), PAR summer (**b**), kd490 spring (**c**), kd490 winter (**d**), and SST winter (**e**). Each row of pie charts represents a coding region annotated with one of the top three GO terms associated with each variable in Fig. 4, grouped by population (SPS, WB, SJF). The colors represent haplotype frequencies, with the most common haplotype colored in red for each term. GO annotations corresponding to the pie chart alphanumeric labels are listed in Supplemental Table S2.

The most common haplotypes detected in SPS for each GO term (Fig. 5a1-3, red) were fixed in WB and present at lower frequencies in SJF (two individuals, Fig. 5a1; 4 individuals in both ab and ac). The most common haplotypes in SJF (Fig. 5a1-3, blue) were also detected at low levels in SPS (in one individual for both 1a and 1c), although they were not present in WB.

In all but two cases (Fig. 5d1, e5), the most common haplotype in SPS was present in all three clusters (Fig 5, red). Differences between the SPS cluster and the other clusters were most pronounced in haplotypes associated with winter SST and spring/winter kd490 (Fig. 5b, d, e). For winter SST, haplotypes unique to SPS were found in three of the five GO groups (Fig. 5b1,4,5; blue), and in one case this haplotype was the most common, present in about 60% of SPS individuals (Fig. 5b5). Notably, for the two remaining GO groups, the SPS-associated haplotype also appeared in two individuals from the WB cluster, both sampled from EB (Fig. 5b2,3, blue; Supplemental Fig. S1). For the kd490-related GO groups, individuals in the WB and SJF clusters consistently shared only a single haplotype, which was also present in SPS (Fig. 5d, e; red). However, between three and ten SPS individuals carried a distinct haplotype for each GO group (Fig. 5d, e2–5; blue), and a third haplotype was detected in two SPS individuals (Fig. 5d1; purple).

#### 3.2.3. Variants which impact gene function

SnpEff identified 36,994 variants with potentially low (LI), moderate (MI), or high impact (HI) on gene function. MI variants were the most common of the three groups, with a total of 20,728 identified across all samples, followed by LI (15,544) and HI (722). Overall, the number of HI and MI variants differed significantly between the three genetic clusters (Fig. 6a,b; p = 4.2e-04, p = 9.5e-04, respectively), though this was not the case with LI (p = 0.061). 89.61% of the HI variants belonged to one of three effect categories: loss of a start codon, loss of a stop codon, or gain of a stop codon (Supplemental Table S3). The remaining HI variants were categorized into two effect categories: splice acceptor site mutations and intron variants, totaling six categories. All MI variants belonged to a single effect category—missense variant—although 0.79% of them were also splice region variants (Supplemental Table S3).

**Figure 6.**
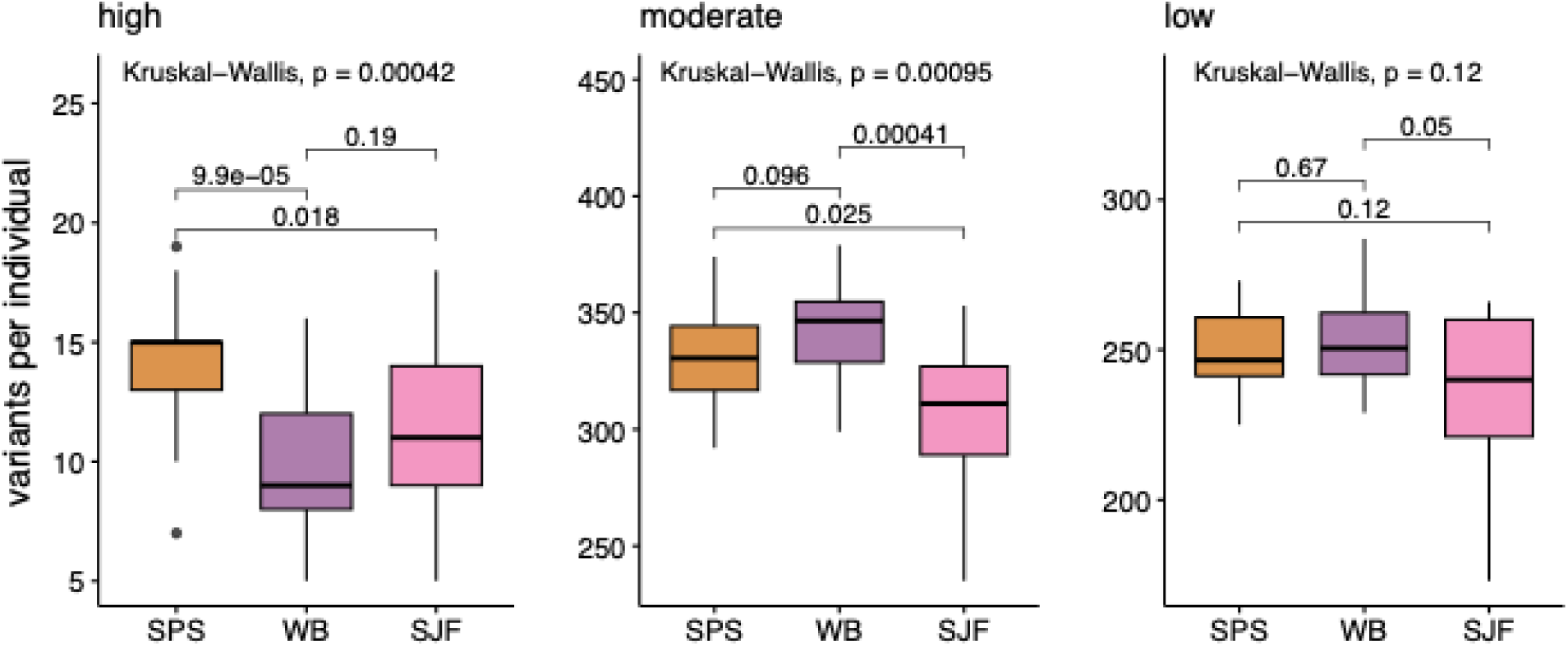
Variants identified by SnpEff per individual per genetic cluster across three populations of *Nereocystis luetkeana* at Puget Sound: South Puget Sound (SPS; organge), Whidbey Basin (WB; purple), and the Strait of Juan de Fuca (SJF; pink). The y-axis represents the number of variants per individual, and the x-axis represents the individual’s population. Each panel represents low (**a**), moderate (**b**), and high-impact variants (**c**). Global p-values (Kruskal-Wallis test) are displayed above the plot, while pairwise comparisons (Wilcoxon test) are denoted above the brackets between the boxes.

Individuals carried an average of 11.46 HI variants. Between the three clusters, SPS individuals had significantly more HI variants on average than WB (p = 9.9e-05, Fig. 6c) and SJF (p = 0.018, Fig. 6c), with 13.94 HI variants per individual on average (ranging between 7-19).

Although HI variants were more common among WB individuals than SJF (𝑥 = 9.96 and 11.29, respectively), this difference was not significant. MI variants were least common in SJF individuals (𝑥 = 306.82) compared to WB (𝑥 = 341.36 p = 0.00041) and SPS (𝑥 = 330.78, p = 0.025), however, there was no significant difference between SPS and WB (p = 0.096). On average, individuals carried 246.73 LI variants, with no significant differences observed between the clusters.

Closer examination revealed that all 722 HI variants were restricted to just 37 loci, spread across 21 scaffolds (Fig.7a, top). None of the loci were unique to a single individual, nor did any individual have HI variants at all 37. On average, each locus was shared between 19.51 individuals. While some were relatively uncommon (fewer than ten individuals shared 14 loci, each), others were quite prevalent: 19 were shared between one quarter of individuals, seven were shared by half, and one was shared by 90% of all individuals (Supplemental Table S4).

When comparing genetic clusters, variants unique to a single group were rare: only two loci were unique to a specific cluster, both of which occurred in SJF. Instead, most of these variants (25) were found in all three clusters.

However, when comparing individuals within a site, fixed HI variants were more common, occurring in 15 of the 19 sampled populations (Fig 7, bottom), excluding Foulweather Bluff (FW) which had only a single individual sampled. Regardless of genetic cluster, each site had at least one HI variant shared between all individuals (Fig 7, bottom). The southernmost site, Squaxin Island (SQ) had the greatest number of fixed HI variants (15), followed by Day Island (DI, twelve) and Salmon Beach (SB, ten), all in SPS. Unexpectedly, Morris Creek (MC) from the SJF cluster, had the same number of fixed HI variants as the Point Vashon (PV) site in SPS (eight). All WB sites had fixed HI variants as well, but these were fewer in number than those in SPS, ranging from one to four HI variants per site. The number of fixed HI variants in SJF was similar: other than MC, the rest ranged between two and four per site.

**Figure 7.**
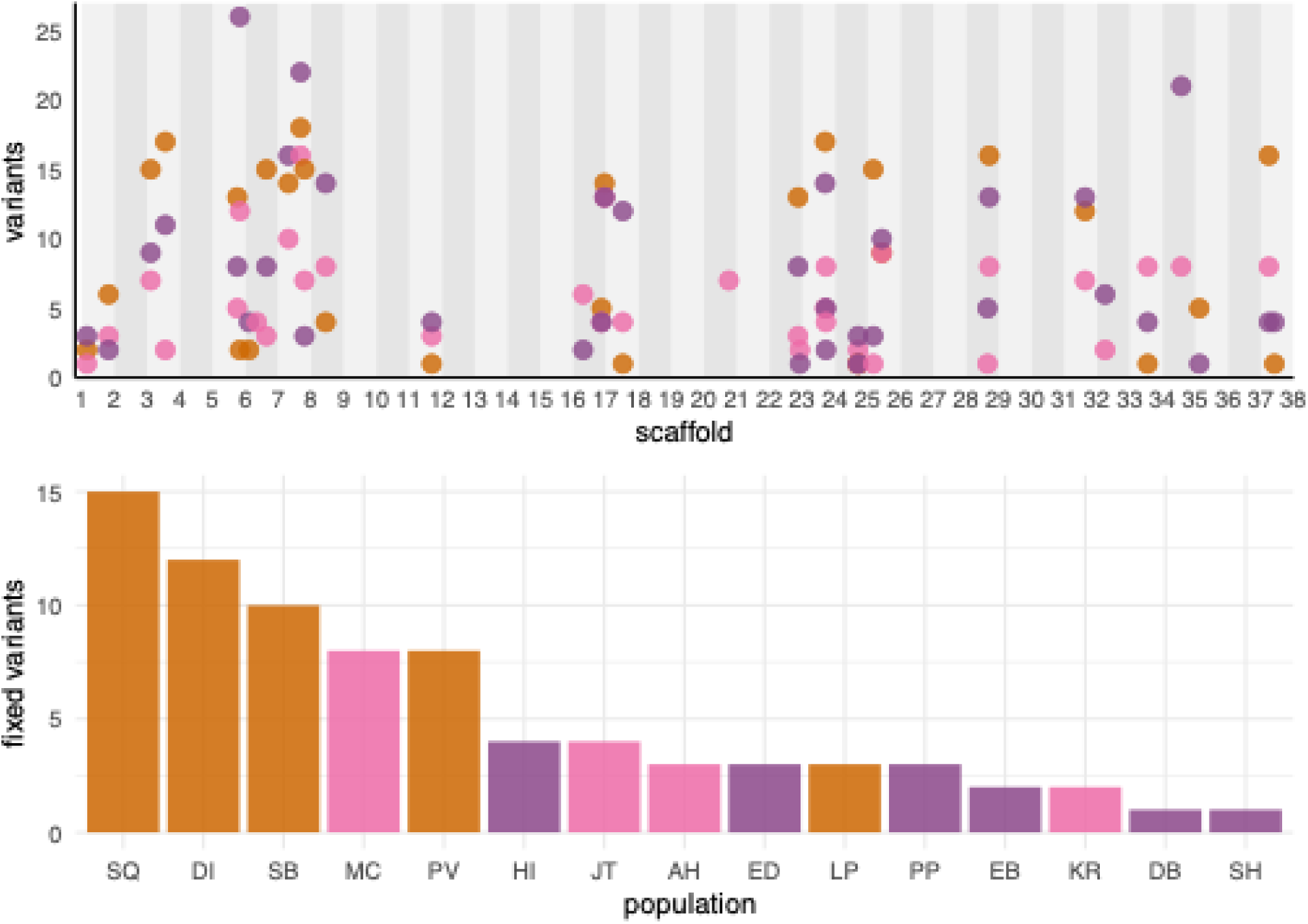
Individual and population HI variants across three populations of *Nereocystis luetkeana* at Puget Sound: Puget Sound (SPS; organge), Whidbey Basin (WB; purple), and the Strait of Juan de Fuca (SJF; pink). **Top:** Position of HI variants across scaffolds (x-axis). Points represent genetic clusters and their height (y-axis) represents the number of individuals that carry a variant at that locus. **Bottom:** HI variants which are present in every individual at a sampled site (x-axis). The height of the bar (y-axis) is the number of HI variants shared by all individuals within that population.

## 4. Discussion

### 4.1. Summary

The primary objective of genetic rescue (GR) is to increase the genetic diversity of a recipient population while minimizing the risk of outbreeding depression. For GR to be successful in the SPS bull kelp populations, selecting an appropriate donor population is a critical first step; it must account not only for overall genetic diversity but also for the compatibility of adaptive genetic variation. Despite low diversity and the prevalence of deleterious alleles, our results suggest that local adaptation is present in Puget Sound bull kelp populations, driven by five of the tested environmental variables. The association of distinct haplotypes with environmental gradients suggests functional variation across sites, indicating that local adaptation in this region may be more prevalent than previously recognized. While these conclusions require experimental validation, they highlight the importance of considering adaptive variation in donor selection.

Although SJF emerges as a promising donor candidate based on its high genetic diversity, the markedly different environmental conditions in this region and the corresponding genetic divergence likely make it a poor match for SPS. This mismatch is particularly evident with respect to salinity, which correlates strongly with environmentally associated genetics in the SJF cluster (Fig. 3b) and is reflected in its distinct functionally significant haplotypes (Fig. 3a). In contrast, WB populations, though less diverse than SJF, may be more suitable because they still offer greater genetic diversity than SPS and also share similar environmentally associated genetics (Fig. 3). A population from the WB cluster would likely be a more suitable candidate. Although less diverse than SJF, it still exceeds the diversity of SPS and shares similar adaptive variation.

### 4.2 Contextualizing functional enrichment results

Salinity showed the strongest correlation with environmentally associated genetic variation (Fig. 2a). This aligns with previous studies that have identified salinity as a key stressor in kelps, with reduced levels known to disrupt growth, photosynthesis, and reproduction (Buschmann et al., <u>2004</u>; Diehl et al., <u>2020</u>; Drakard et al., <u>2025</u>; Lind and Konar, <u>2017</u>; Monteiro et al., <u>2020</u>; Miyashita, <u>2007</u>; Vettori et al., <u>2020</u>). Salinity varies most at the boundary between SJF and WB, mainly due to freshwater inflow into WB from rivers, watersheds, outfalls, and snowmelt (Yang and Khangaonkar, <u>2010</u>). This contrast is mirrored when considering functionally significant genes, where the most common SJF haplotype was either rare or absent in the other clusters.

Furthermore, salinity-associated GO terms were primarily related to the maintenance of cellular homeostasis, such as mitigating osmotic, metabolic, and/or oxidative stress. In plants and algae, purine metabolism is dynamically regulated in response to changes in salinity (Suzuki-Yamamoto et al., <u>2006</u>; Wei et al., <u>2017</u>; Zhang et al., <u>2011</u>; Zheng et al., <u>2024</u>). Furthermore, in marine algae, taurine dioxygenase and histidine-tRNA ligase serve as osmoregulators linked to salinity stress tolerance, and glutathione peroxidase is known to combat salinity-induced oxidative stress (Dabravolski & Isayenkov, <u>2024</u>; Islam et al., <u>2015</u>; Jahnke & White, <u>2003</u>; Judy et al., <u>2024</u>; Kumar et al., <u>2010</u>; Pinseel et al., <u>2022</u>; Rani, <u>2007</u>; Yihua et al., <u>2024</u>).

While salinity was the overall greatest predictor, SST winter is likely the greatest driver of natural selection in SPS (Fig. 3b, blue). SST winter was correlated mainly with a site’s distance from the ocean, with the coolest temperatures in SJF and warmest in SPS. Notably, it is important not to conflate adaptation to winter temperatures with adaptation to cold. Without experimental validation, it is unclear whether the association reflects selection for low temperatures generally or for the comparatively warm winters of SPS. Haplotype analyses of functionally significant genes revealed that SPS individuals possessed sequences that were absent (Fig. 5b1, b4, b5) or rare (Fig. 5b2, b3) in other clusters. Similar associations have been reported in other studies which broadly fell into two GO terms helicases and the pyrimidine biosynthesis pathway, which are both linked to cold response in plants (Bellin et al., <u>2023</u>; Gong et al., <u>2002</u>; 2005; Lei et al., <u>2022</u>; Mohapatra et al., <u>2023</u>; Vos et al., <u>2007</u>; Zhang & Tak, <u>2020</u>)

Summer light availability (PAR summer) varied greatly across SJF cluster range, which contained the sites with both the highest and lowest values. Light availability was similarly low around Whidbey Island compared to SPS and western SJF. This may explain the two haplotypes unique to SJF (Fig. 5c1,3), as well as the three others shared only with WB (Fig. 5c2). Nearly half of the PAR summer GO terms were linked to metabolites produced by plants and algae to counter photo-inhibition and oxidative damage under high light stress (trehalose and methionine; (Dong et al., <u>2016</u>; Kumar et al., <u>2017</u>; Levine et al., <u>2008</u>; Rathod et al., <u>2016</u>; Rathod et al., <u>2023</u>; Sadak et al., <u>2019</u>; Zhang et al., <u>2023</u>). Most of the remaining PAR summer GO terms were related to various motor protein functions often associated with light-induced chloroplast movement, which enables photosynthetic organisms to optimize photosynthesis in low-light conditions and prevent photodamage from excessive light (Masamitsu, <u>2016</u>; Suetsugu & Wada, <u>2007</u>; Suetsugu et al., <u>2016</u>; Wada et al., <u>2003</u>).

Turbidity (kd490) varied substantially across sites and seasons, with the lowest levels consistently observed in SJF, with peak spring values in WB and peak winter values in SPS. Turbidity is associated with a variety of stressors, including reduced light, eutrophication, sedimentation, or pollution. However, as kd490 does not identify particles in the water column, identifying the precise stressor is beyond the scope of this study. An added layer of complexity is that the types of particles could vary between sites, resulting in a wide range of GO terms with little consistency. However, four of the six terms were linked by a common function in marine algae: resistance to metal stress (Li et al., <u>2013</u>; Qiao et al., <u>2021</u>; Qu et al., <u>2023</u>; Sharma et al., <u>2019</u>). Water quality assessments support this interpretation: several SPS sites with severe metal pollution have been identified, including contaminants such as arsenic, chromium, mercury, lead, nickel, and zinc (Washington State Department of Ecology, 303(d), 1996-2018). Some of the kd490-associated GO terms are linked to additional functions: nitric oxide is linked to microplastic and pesticide defense (Li et al., <u>2013</u>; Wang et al., <u>2025</u>), while inositol oxygenase is linked to wastewater tolerance (Qiao et al., <u>2021</u>). This pattern indicates that the kd490-associated GO terms—and the unique haplotypes found in SPS—are likely linked to adaptation to metal stress, a factor not present in WB, where such contaminants are largely absent and no distinct haplotypes were observed (Washington State Department of Ecology, 303(d), 1996-2018; U.S. Department of Interior & U.S. Geological Survey, <u>2018</u>).

### 4.3. Future directions

Although seascape genomics has many advantages, it is not without limitations. One key issue is that the environmental variables that are most strongly associated with adaptive variation in SPS (winter SST and turbidity, Fig. 3b) are linked to functionally significant haplotypes that are largely unique to SPS, with only two occurring outside the region (Fig. 5b2, b3). Without experimental validation, such as a reciprocal transplant between WB and SPS, it is unclear whether this disparity would increase the risk of outbreeding depression. Furthermore, since all individuals are preserved in our GenBank, laboratory-based gene expression experiments could help determine whether the identified genes are truly functionally significant. Such experiments would also allow us to test which WB genotypes are best suited to SPS environmental conditions and disentangle the specific selective pressures associated with turbidity—such as low light, metal toxicity, or pollution—that may be driving local adaptation. However, whether WB genetic diversity is sufficient to serve as a donor for GR remains an open question and has yet to be tested. Low diversity in SPS has limited the effectiveness of natural selection in purging deleterious alleles, leading to an increased genetic load (Bemmels et al., <u>2025</u>). Theoretical models have shown that crossing two inbred populations can still alleviate inbreeding depression, provided that each carries different deleterious alleles that can be masked by alleles from the other (Charlesworth & Charlesworth, <u>1999</u>; Lynch, <u>1999</u>; Tallmon et al., <u>2004</u>). While experimental validation is needed to determine whether deleterious alleles from one population can indeed be masked by the other, we can estimate their prevalence based on our SnpEff results. Our results show that HI variants (which cause loss of function) are most common in SPS (Fig. 6, left); however, unlike SPS, relatively few of these variants are fixed in WB populations (Fig. 7). Thus, it is possible to select WB individuals that both share adaptive genetic variation with SPS and lack its fixed deleterious variants. Ultimately, these results underscore the role of local adaptation, support the suitability of a population from WB as a donor, and lay the groundwork for future research and conservation efforts.

## Acknowledgements

This study was supported by the grant from Washington Sea Grant/University of Washington (Award No. UWSC13569) awarded to Jodie Toft, Hilary Hayford, and Filipe Alberto. We would like to thank Puget Sound Restoration Fund (PSRF) for consulting on this project and collecting samples for the biobank, especially Hilary Hayford, Brian Allen, Jessi Florendo, Sofia O’Connell, Max Calloway, and Kari Inch. We would also like to thank NOAA’s Manchester Research Station for hosting the PSRF kelp laboratory. The biobank at the University of Wisconsin Milwaukee is maintained by Gabriel Montecinos, Shanice Piango, and Begona Ramirez Ibaceta. Sequencing was performed by Great Lakes Genomics Center, and the bioinformatics analysis was accomplished using computational resources provided by the Texas Advanced Computer Center.

## Data availability

Raw reads used in this study can be accessed under the NCBI BioProject accession PRJNA1335537, and the annotated reference genome is publicly available through Joint Genome Institute Phycocosm portal (https://phycocosm.jgi.doe.gov/Nerluet1/Nerluet1.home.html). All scripts used in this analysis can be found on GitHub at https://github.com/evelynabbott/bullkelp_WGS.

## Funding statement

This project was supported by the grant from the Washington Sea Grant/University of Washington (Award No. UWSC13569) awarded to Jodie Toft, Hilary Hayford, and Filipe Alberto.

## Conflict of interest

The authors declare no conflict of interest.

## Notes

https://www.ncbi.nlm.nih.gov/bioproject/?term=PRJNA1335537

### Competing Interest Statement

The authors have declared no competing interest.

