## Supplementary figures and images for "Evidence of local adaptation in Puget Sound’s threatened bull kelp (*Nereocystis luetkeana*) populations"

### Supplemental Fig. S1

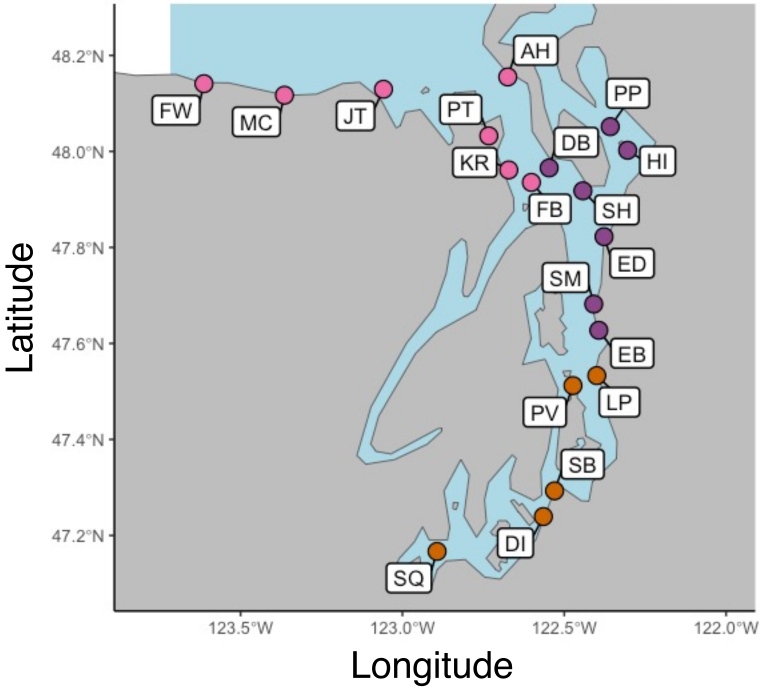

### Supplemental Fig. S2

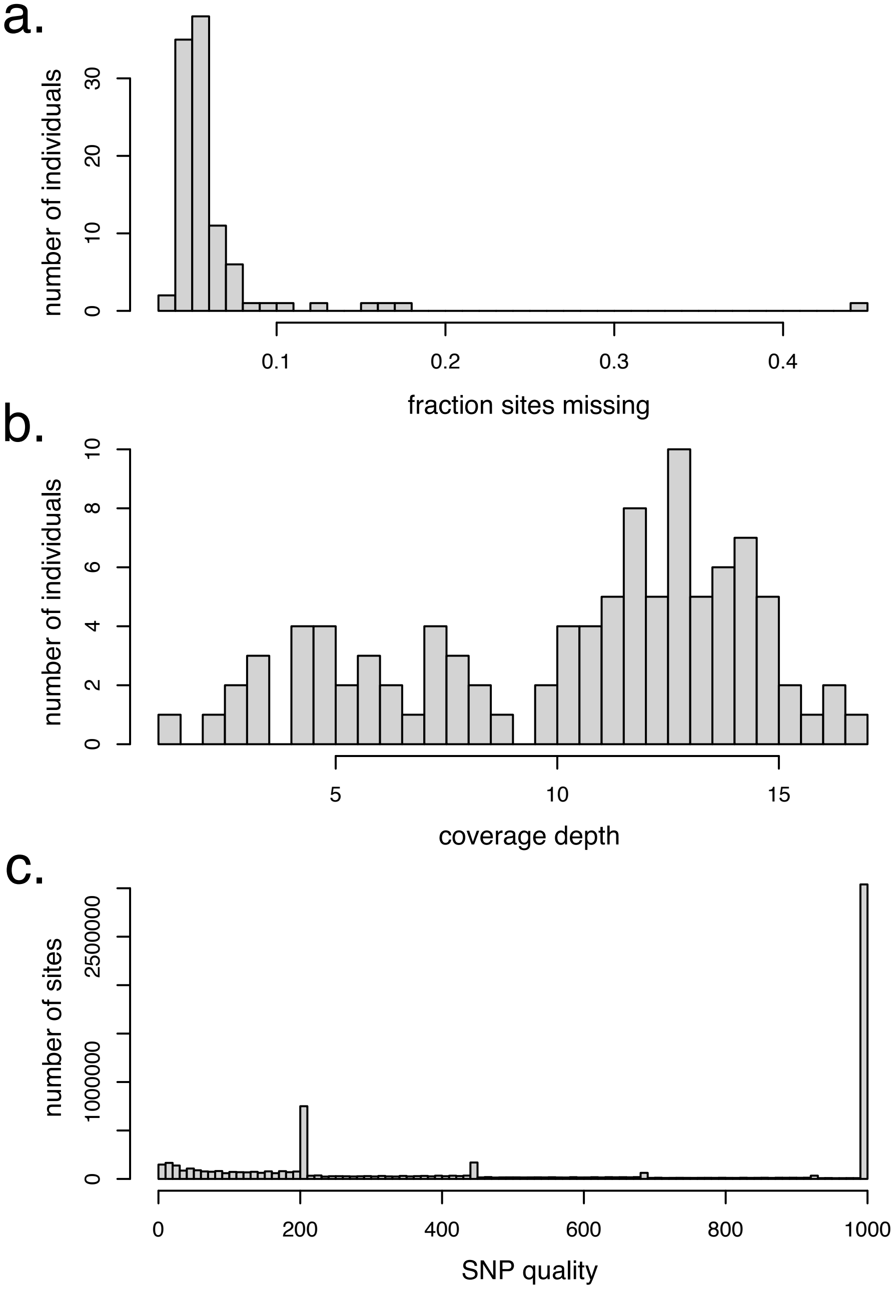
